# Post-ictal generalized EEG suppression and seizure-induced mortality are reduced by enhancing dorsal raphe serotonergic neurotransmission

**DOI:** 10.1101/2020.06.28.172460

**Authors:** Alexandra N. Petrucci, Katelyn G. Joyal, Jonathan W. Chou, Rui Li, Kimberly M. Vencer, Gordon F. Buchanan

## Abstract

Sudden unexpected death in epilepsy (SUDEP) is the leading cause of death in patients with refractory epilepsy. A proposed risk marker for SUDEP is the duration of post-ictal generalized EEG suppression (PGES). The mechanisms underlying PGES are unknown. Serotonin (5-HT) has been implicated in SUDEP pathophysiology. Seizures suppress activity of 5-HT neurons in the dorsal raphe nucleus (DRN). We hypothesized that suppression of DRN 5-HT neuron activity contributes to PGES and increasing 5-HT neurotransmission or stimulating the DRN before a seizure would decrease PGES duration. Adult C57BL/6 and *Pet1-Cre* mice received EEG/EMG electrodes, a bipolar stimulating/recording electrode in the right basolateral amygdala, and either a microdialysis guide cannula or an injection of adeno-associated virus (AAV) allowing expression of channelrhodopsin2 plus an optic fiber into the DRN. Systemic application of the selective 5-HT reuptake inhibitor citalopram (20 mg/kg) decreased PGES duration from seizures induced during wake (n = 23) and NREM sleep (n = 13) whereas fluoxetine (20 mg/kg) pretreatment decreased PGES duration following seizures induced from wake (n = 11), but not NREM sleep (n = 9). Focal chemical (n = 6) or optogenetic (n = 8) stimulation of the DRN reduced PGES duration following kindled seizures and reduced morality following maximal electroshock seizures (n = 6) induced during wake. During PGES, animals exhibited immobility and suppression of EEG activity that was reduced by citalopram pretreatment. These results indicate that 5-HT and the DRN may regulate PGES and seizure-induced mortality.

**Highlights:**

- PGES consistently follows seizures induced by amygdala stimulation in amygdala-kindled mice.
- Seizure-induced dysregulation of 5-HT neurotransmission from the dorsal raphe nucleus may contribute to PGES.
- Systemic administration of 5-HT enhancing drugs and stimulation of the DRN reduces PGES duration.
- PGES is associated with post-ictal immobility in kindled mice that can be reduced by pretreatment with citalopram.
- Recovery of EEG frequencies to baseline occurs in a stepwise manner with the lowest frequencies recovering first.

## Introduction

Epilepsy is a highly prevalent neurological disease characterized by recurrent, spontaneous, and often unpredictable seizures (Fisher et al., 2014). It is estimated that one in twenty-six Americans will develop epilepsy during their lifetime (Hesdorffer et al., 2011). Approximately one-third of these patients will not achieve seizure freedom with currently available therapies (Chen et al., 2018; Kwan and Brodie, 2000), putting them at greater risk for sudden unexpected death in epilepsy (SUDEP). SUDEP is the leading cause of death in individuals with uncontrolled seizures (Devinsky et al., 2016) and is second only to stroke in years of potential life lost due to neurological disease (Thurman et al., 2014). While cardiac (Auerbach et al., 2013; Richerson, 2013), respiratory (Richerson, 2013; Richerson and Buchanan, 2011), and arousal impairment (Buchanan, 2019; Richerson, 2013) are involved, the exact etiology of SUDEP is unknown. Serotonin (5-HT) has been implicated in SUDEP (Petrucci et al., 2020) and seizure-induced death (Buchanan et al., 2014; Faingold et al., 2014) due to its role modulating seizures (Bagdy et al., 2007; Petrucci, et al., 2020), breathing (Buchanan et al., 2015; Richerson, 2004), and sleep/wake and arousal (Buchanan, 2019; Purnell et al., 2018). Administration of 5-HT enhancing drugs reduces seizure frequency in epilepsy patients (Ceulemans et al., 2012; Favale et al., 2003; Favale et al., 1995). This is recapitulated in rodent seizure models including genetically epilepsy prone rats (GEPR) (Dailey et al., 1992), EL mice (Imaizumi et al., 1959; Kabuto et al., 1994), and audiogenic seizure prone DBA mice (Tupal and Faingold, 2006).

A potential risk marker for SUDEP is the duration of post-ictal generalized EEG suppression (PGES), a period of low amplitude, low frequency electrographic activity following some seizures (Lhatoo et al., 2010; Okanari et al., 2017). Patients are more likely to be immobile, unresponsive, and require resuscitative measures during PGES (Kuo et al., 2016; Semmelroch et al., 2012). However, PGES remains poorly understood and its origin is unknown. Recently, a study in humans observed that interictal serum 5-HT level is inversely correlated with PGES duration (Murugesan et al., 2018). While serum 5-HT does not directly relate to parenchymal 5-HT, it does support involvement 5-HT or 5-HT producing neurons in PGES. Here we examine the mechanism underlying the effects of 5-HT on PGES.

There are two major populations of 5-HT neurons within the brain: a rostral group and a caudal group (Dahlstroem and Fuxe, 1964). The dorsal raphe nucleus (DRN) is a major component of the rostral group (Hornung, 2010). 5-HT neurons within the DRN send projections to regions associated with sleep/wake regulation (Monti, 2010; Steininger et al., 1997) and arousal (Vertes and Kocsis, 1994). Moreover, SUDEP is thought to be more likely to occur during sleep (Lamberts et al., 2012) and it is known that DRN 5-HT neurons fire fastest during wake, slow down during non-rapid eye movement (NREM) sleep, and are nearly quiescent during REM sleep (McGinty et al., 1973). DRN firing is also depressed by seizures (Zhan et al., 2016). Therefore, we hypothesized that seizure-induced dysregulation of the DRN 5-HT network may contribute to PGES and that increasing 5-HT tone via 5-HT enhancing drugs or DRN stimulation would reduce PGES and seizure-induced mortality.

There are very few animal studies examining PGES. Here we establish amygdala kindling as a reliable model of PGES and examine the role of 5-HT on behavior and EEG characteristics during PGES. We tested our hypothesis in amygdala kindled mice by increasing 5-HT tone via (1) systemic administration of selective serotonin reuptake inhibitors (SSRIs), (2) focal stimulation of the DRN with acidosis, and (3) optogenetic stimulation of the DRN prior to seizure induction. Because SUDEP likely occurs more often during sleep and 5-HT is regulated in a sleep/wake dependent manner (Lamberts, et al., 2012; McGinty and Harper, 1976; Purnell, et al., 2018), when possible seizures were induced during different sleep/wake states.

## Experimental Procedures and Methods

### Animals

All procedures and experiments were approved by and performed in compliance with the Institutional Animal Care and Use Committee at the University of Iowa Carver College of Medicine. All experiments complied with the National Research Council’s guide for the care and use of laboratory animals (2011). Animal pain and distress was minimized during experimental procedures. Adult (6-20 week) male and female C57BL/6J and *Pet1-Cre* mice were used for these studies. *Pet1-Cre* mice (B6.Cg-Tg[Fev-cre]1Esd/J) (Scott et al., 2005) were originally obtained from Jackson Labs (Bar Harbor, ME), subsequently crossed to C57BL/6J, and maintained in our animal care facilities at University of Iowa. *Pet1-Cre* mice express Cre-recombinase exclusively in *Pet1* expressing neurons, enabling Cre-dependent targeting to 5-HT neurons (Hendricks et al., 1999). Mice were housed in 12:12 light:dark cycle (6:00 AM to 6:00 PM or 12:00 PM to 12:00 AM) with food and water available *ad libitum*. Experiments were performed during the same circadian time window between 2-6 hrs after the start of the light phase. Experiments were spaced at least two days apart to avoid confound of repeated stimulations.

### Surgical procedures

#### General

Surgical procedures were performed under anesthesia (isoflurane gas 2-5% induction; 0.5-2% maintenance) using aseptic technique. Surgical plane of anesthesia was verified via toe pinch every 15 min and anesthesia adjusted as needed. Mice were secured into a stereotaxic apparatus (51730; Stoelting Co.; Wood Dale, IL). The heads were shaved, cleaned with betadine and 70% ethanol, and a ∼2.0 cm longitudinal incision was made in the scalp. Throughout surgery and initial recovery, a heating pad was used to regulate body temperature (Physiosuite; Kent Scientific; Torrington, CT). For pain management, animals received meloxicam (2.0 mg/kg; *i*.*p*.) pre-operatively and at 24 and 48 hrs post-operatively. Animals recovered for at least seven days before being studied.

#### EEG/EMG headmount implantation and bipolar stimulating/recording electrode implantation

After being prepared as above, six holes were drilled into the skull using a 0.5 mm drill bit. The holes were positioned as follows: two 2 mm ±2 mm anterior to bregma, two 2 mm ±2 mm anterior to lambda, one overlying the right basolateral amygdala (in mm from bregma with skull level: AP: −1.3, ML: −2.8, DV: −4.7), and one equidistant between the left holes. Unless specified, holes were made approximately 1.5 mm lateral to midline. Small screws (1.0 mm thread, 4.1 mm length, 0.8 mm shaft diameter; #80404; Vigor Optical; Carlstadt, NJ) soldered to stainless steel wires (793000; A-M Systems; Carlsborg, WA) were threaded into four holes to serve as EEG electrodes. A ground screw without wire was threaded in the hole between the left EEG electrodes. Then a polyimide coated, bipolar twisted pair stimulating/recording electrode (MS333-3-BIU-SPC; Plastics One, Inc; Roanoke, VA) with the distal 0.5 mm of insulation removed was implanted into the right basolateral amygdala. The implant was secured with thick cyanoacrylate (PT-33; Pacer Technology – ZAP; Ontario, CA) and dental acrylic before the stainless steel ground wire was soldered to the ground screw.

After insertion of additional implants as described below, the wires attached to the EEG screws were twisted and soldered to wires protruding from a six-pin socket (853-41-006-30-001000; Millmax Co.; Oyster Bay, NY). EMG electrodes on the headmount were inserted bilaterally into the nuchal muscles. Dental acrylic (Lang Dental; Wheeling, IL) was applied over the EEG wires to increase stability and decrease electrical noise. The overlying skin was closed with 5-0 nylon suture (668G; Ethilon, Inc.; Somerville, NJ).

#### Microdialysis cannula implantation

C57BL6/J mice undergoing chemical stimulation of the DRN were implanted with a guide cannula (CMAP000138 CMA7; CMA, Inc; Kista, Sweden) into the DRN (AP: −4.6, ML: 0, DV: −3.0) during the EEG/EMG headmount and amygdala electrode surgery (Smith et al., 2018). Guide cannula were secured with thick cyanoacrylate and dental acrylic. Identical metal-free cannula (CMA8010773; CMA, Inc) were implanted for MES experiments to prevent electrical propagation along metal components.

#### Adeno-associated virus administration and optic fiber insertion

*Pet1-Cre* mice and *WT* littermates were instrumented for EEG/EMG recording and amygdala kindling as described. After insertion of epidural screws, an adeno-associated virus (AAV) containing Channelrhodopsin2 (ChR2; rAAV5-EF1a-DIO-hChR2[H134R]-mCherry; University of North Carolina Gene Therapy Center Vector Core; Chapel Hill, NC; Titer 1.8×10^12^ vg/ml), was injected into the DRN using either a 2 µl Hamilton syringe (6545901; Hamilton Company; Reno, NV) or an autoinjector (70-4507; Harvard Apparatus; Holliston, MA) at a rate of 0.5 µl over 10 min and allowed to diffuse for another 10 min. A polyimide optical fiber (MFC_200/240-0.22_4mm_ZF1.25_FLT; Doric Lenses; Québec, Canada) was then implanted into the DRN and secured with thick cyanoacrylate and dental acrylic. Animals receiving viral vectors were not tested for at least 28 days following surgery to allow transfection.

### Seizure models

#### Amygdala kindling

A fast-kindling paradigm was utilized for amygdala kindling as described previously (Santoro et al., 2010). An EEG/EMG preamplifier (8202-SL; Pinnacle Technology; Lawrence, KS) and a 3-channel electrode cable (335-340/3; Plastics One) were attached to the animal’s headmount and connected to a data conditioning amplifier (LP511AC; AstroNova; West Warwick, RI), a stimulating/recording switch (SRS-13C0113G; AstroNova), and a pulse stimulator (Model 2100; A-M Systems). To determine the amygdala afterdischarge threshold, 20 µA incremental currents were administered every two minutes until after-discharges were recorded. The threshold stimulus was applied twice daily at least 1 hr apart (80-500 µA range, 1 s train of 1 ms biphasic square wave pulses at 60 Hz) until the animal had three consecutive Racine grade 4 generalized seizures as rated on a modified Racine scale as follows: 1, behavioral arrest; 2, automatisms, drooling, whisker twitches, body jerks; 3, unilateral limb myoclonus; 4, bilateral myoclonus, loss of posture, wild running; 5, status epilepticus; 6, death (Buchanan, et al., 2014; Racine, 1972).

#### Maximal electroshock seizure induction

C57BL/6J mice (control group: n = 6, 3 male, 3 female; acidosis group: n = 6, 2 male, 4 female) were acclimated to the recording apparatus and ear-clip electrodes (modified, toothless, stainless steel alligator clips coated with saline-moistened gauze) for at least 1 hr per day on two consecutive days. On trial days the ear-clip electrodes were attached to the mice and plugged into a Rodent Shocker pulse generator (D-79232; Harvard Apparatus). After a 30 min habituation period, a single 50 mA (0.2 s, 60 Hz sine wave pulses) stimulus was administered during wake to induce a maximal hindlimb extension seizure (Buchanan, et al., 2014; Hajek and Buchanan, 2016; Purnell et al., 2017). Mortality, seizure duration, and extension-to-flexion ratios (E/F ratios) were determined post-hoc using EEG/EMG monitoring and video assessments. Higher E/F ratios correlate with widespread propagation of epileptiform activity and are used as an indicator of seizure severity (Anderson et al., 1986). E/F ratios were calculated post-hoc via video analysis to measure the duration of hindlimb extension (> 90°) divided by the duration of hindlimb flexion (≤ 90°).

### Experimental procedures

#### Systemic Drug Administration

Adult male and female C57BL/6J mice were instrumented for EEG/EMG recording and amygdala kindling. After surgical recovery and completion of kindling, mice received an *i*.*p*. injection of citalopram (20 mg/kg; wake: n = 23, male = 12, female = 11; NREM: n = 13, male = 8, female = 5), fluoxetine (10 mg/kg; wake: n = 11, male = 6, female = 5; NREM: n = 9, male = 4, female = 5), or saline (wake: n = 30, male = 15, female = 15; NREM: n = 19, male = 12, female = 7) 30-60 min prior to induction of an seizure during wake or NREM sleep. Citalopram hydrobromide and fluoxetine hydrochloride were obtained from Tocris Biosciences through Bio-Techne (Minneapolis, MN). Dosages were based upon the lab’s previous experience with the drugs and the primary literature (Buchanan, et al., 2014). Drugs were administered in consistent volumes (0.15 - 0.2 ml).

#### Reverse microdialysis for delivery of normal and acidified aCSF

During reverse microdialysis trials mice were attached to cabling for EEG/EMG recording and the dummy cannula was replaced with either a metal dialysis probe for kindling experiments (pores <6 kDa, tip length 1 mm, tip diameter 240 μm; CMAP000082; CMA7; CMA Inc; n = 6, 2 male, 4 female) or a metal-free dialysis probe for the MES experiments (CMA8010771; CMA7; CMA Inc.; pores <6 kDa; tip length 1 mm; tip diameter 240 μm; control group: n = 6, 3 male, 3 female; acidosis group: n = 6, 2 male, 4 female).

Normal artificial cerebrospinal fluid (aCSF) (bubbled with 5% CO_2_ / 21% O_2_ / balance N_2_; pH 7.4) was dialyzed using a syringe pump (45 μl/min; 70-3005, Harvard Apparatus) throughout the 20 min habituation period. The dialysate was then changed to aCSF bubbled with 25% CO_2_ / 21% O_2_ / balance N_2_ (pH 6.8) or another syringe with normal aCSF for 10 min prior to seizure induction as described (Dias et al., 2008; Li et al., 2013; Smith, et al., 2018). Gas tanks were obtained from Praxair (Cedar Rapids, IA). The composition of the aCSF (Smith, et al., 2018) was as follows (in mM): 152 Na^+^, 3.0 K^+^, 2.1 Mg^2+^, 2.2 Ca^2+^, 131 Cl^−^, and 26 HCO_3_ ^−^. The Ca^2+^ was added after the aCSF was warmed to 37 °C and equilibrated with CO_2_.

#### Optogenetic manipulation of 5-HT neurons

*Pet1-Cre* mice and their *WT* counterparts were instrumented for EEG/EMG and amygdala kindling. Mice were injected with AAV-ChR2 (*Pet1-Cre:* n = 8, 3 male, 5 female; *WT*: n = 7, 3 male, 4 female) and instrumented with an optic fiber (4 mm, 240 µm diameter, 0.22 NA; MFC_200/240-0.22_4mm_ZF1.25_FLT; Doric Lenses) in the DRN. On the day of trials, light power (∼12.5 mW to account for 25% loss) was tested with a digital optical power and energy meter (P0014083; Thor Labs; Dachau, Germany). Mice were placed into the recording chamber and plugged into the EEG/EMG lead, the amygdala electrode, and an optical fiber patch cord (MFC_200/240/900-0.22_0.6m_FC-ZF1.25[F]; Doric Lenses). Baseline EEG was recorded for ∼5 min. Animals received light stimulation (473 nm, 4 Hz, 10 mW, 2% duty, 10 s ON / 10 s OFF; Opto Engine, LLC.; Midvale, UT) or no light stimulation 10 min prior to seizure induction during wake. Laser vs no laser trials were randomized to account for number of seizure inductions. Optogenetic manipulation prevented animals from falling into a deep sleep, precluding trials during NREM. Wake was confirmed offline by EEG/EMG analysis and video.

#### Post-ictal immobility determination

Post-ictal immobility was quantified following kindled seizures in C57BL/6J mice through video analysis by two reviewers (A.N.P and K.M.V). Each reviewer recorded latency to the first twitch, paw movement (unilateral, no change in body position), and body movement (bilateral limb movement, body position change, preening) after seizure termination. We utilized an interrater reliability threshold of ≥90%. Both reviewers re-evaluated video for data points with <90% interrater reliability until consensus was reached. Reviewer scores were then averaged to produce final quantification. Post-ictal immobility was assessed in mice that received *i*.*p*. citalopram (wake: n = 23, 12 male,11 female; NREM: n = 13, 8 male, 5 female), fluoxetine (wake: n = 11, 6 male, 5 female; NREM: n = 10, 5 male, 5 female), or saline (n = 55; 32 male, 23 female) 30-60 min prior seizure induction. Video was not analyzed in mice whose limbs were obscured.

### EEG data collection and analysis

#### EEG/EMG data acquisition

Mice were attached to a preamplifier (8202-SL; Pinnacle Technology) connected a commutator (8204; Pinnacle Technology) and digital conditioning amplifier (Model 440 Instrumentation Amplifier; Brownlee Precision; San Jose, CA). The EEG signals were amplified 50,000x and band-pass filtered from 0.3 to 200 Hz. The EMG signals were amplified 50,000x and band-pass filtered from 10 to 1000 Hz. Data were digitized with an analog-to-digital converter (PCI-6221 or NI-USB-6009; National Instruments; Austin, TX) at 1000 Hz. The data were then compiled using custom MATLAB software.

#### Sleep state determination

Sleep state was determined on-line and verified post-hoc using a standard approach based on the EEG/EMG frequency characteristics as described previously (Buchanan and Richerson, 2010; Buchanan, et al., 2015; Franken et al., 1998; Smith, et al., 2018): Wake: low-amplitude, high-frequency (7–13 Hz) EEG with high EMG power; NREM sleep: high-amplitude, low-frequency (0.5–4 Hz) EEG with moderate to low EMG power and lack of voluntary motor activity. Video was recorded to assist with vigilance state determination and for verification post-hoc. It is not possible to be blind to vigilance state on-line, but offline analysis was performed blind to vigilance state.

#### EEG frequency band activity analysis

EEG data for frequency analysis was extracted from mice in the aforementioned experiments (1-12 Hz: wake: n = 31, 17 male, 14 female; NREM: n = 19, 12 male, 7 female; 12-30 Hz: wake: n = 26, 12 male, 11 female; NREM: n = 13, 8 male, 4 female). The frequency bands of interest were: delta (δ), 1 – 4 Hz; theta (θ), 4 – 8 Hz; alpha (α), 8 – 12 Hz; beta (β), 12 – 30 Hz. To determine the latency to resumption of baseline activity in certain frequency bands, the power of each band was compared with the lower limit of its 95% confidence interval during baseline. EEG signals were first trimmed to a total length of 500 s (1000 Hz sample rate; 200 s before and 300 s after the seizure ended). The time-frequency information was then extracted via variable wavelet cycles using EEGLAB toolbox. The baseline 95% confidence interval for frequency bands was calculated using the average power across the 200 s pre-seizure period. Post-ictal time-frequency power was smoothed using a moving 5 s window. These results were compared to the corresponding lower limit of 95% confidence interval. A frequency band was considered recovered when power across the selected frequencies stayed above the lower limit for ≥ 5 s.

#### PGES determination

There are discrepancies even amongst experts in PGES identification (Theeranaew et al., 2018). PGES duration was assessed automatically using a custom MATLAB (Mathworks) script and event-related spectral perturbation (ERSP) time-frequency spectral decompositions of the preictal, ictal, and post-ictal period in the MATLAB EEGlab toolbox (Swartz Center for Computational Neuroscience, San Diego, CA). Variable wavelet cycles were used to extract power-frequency parameters from the EEG signal. PGES duration was calculated in a moving window (5 s). The mean EEG power of pre-ictal baseline was subtracted from the mean of the moving window until post-ictal EEG power exceeded 3 standard deviations of baseline EEG power for at least 5 s. ERSP power values are expressed in decibel (dBs) as a measure of relative intensity compared to a baseline pre-ictal period (100s). See Roach and Mathalon (2008) for review (Roach and Mathalon, 2008). Accuracy of the analysis was verified by examining the duration of negative dB values within the spectral plots.

### Tissue processing and statistics

#### Intracardiac perfusion

At the completion of experiments, animals were anesthetized with an overdose of ketamine/xylazine (*i*.*p*., 50–75 mg/kg; 5–7.5 mg/kg) prior to perfusion. Depth of anesthesia was confirmed by toe pinch prior to making an abdominal incision. The diaphragm was cut to expose the heart, the right atrium was punctured, and a 25 gauge needle was inserted into the left ventricle. Chilled (4°C) phosphate buffered saline (PBS) was perfused (∼17 ml/min) until blood was flushed from the circulation. Then chilled 4% paraformaldehyde (PFA) was perfused to fix the tissue. The brains were extracted and placed into 4% PFA at 4°C for 24 hrs. Brains were then cryoprotected in 30% sucrose until the tissue was saturated enough to sink in the solution. Brains were then embedded in Tissue-Tek optimum cutting temperature compound (VWR International; Radnor, PA) and stored at −80°C until they were sectioned on a cryostat (30 μm; Leica Biosystems; Buffalo Grove, IL).

#### Histology

Cell bodies were visualized with Nissl staining to verify placement of cannula and electrodes. Verification of AAV transfection and location was confirmed via immunohistochemistry. Primary and secondary antibodies were diluted with 1x PBS-T containing 1:50 normal horse serum and 0.05% NaN_3_ to permeabilize tissue membranes, block nonspecific binding, and prevent bacterial contamination. Tissue was washed 4-6 times in 1x PBS prior to being bathed in primary antibodies overnight ([mouse anti-TpOH; 1:30,000; MAB5278; EMD Millipore; Burlington, MA], [rabbit anti-DsRed; 1:1000; 632496; Takara Bio Inc.; Kusatsu, Japan], or [chicken anti-GFP; 1:3000; A10262; Invitrogen; Carlsbad, CA]). Tissue was then washed in 1x PBS 4-6 times before a 2 hr exposure to secondary antibodies (Alexa 488 donkey anti-mouse, 1:500, 715545150; Cy3 donkey anti-rabbit, 1:500, 712165153; Jackson Immunoresearch; West Grove, PA]. Anatomic landmarks were identified with the aid of a mouse brain atlas (Franklin and Paxinos, 2013). Fluorescent images were obtained utilizing confocal microscopy and post-processed in OlyVIA software (Olympus Life Science; Center Valley, PA).

#### Statistical analyses

Normality of the data was assessed via a Shapiro-Wilk normality test. Statistical significance was computed using a Mann-Whitney U test unless otherwise specified. Data were analyzed using Graphpad Prism (San Diego, CA). Data were represented as mean ± SEM. Threshold for statistical significance was set at *p* < 0.05 for all comparisons. Post-hoc power analyses were performed using G*Power (open source, Heinrich Heine University Düsseldorf) and to confirm ≥0.80 β power for experiments.

## Results

### Induction of seizures in amygdala kindled mice consistently produced PGES

Electrical stimulation of the amygdala in kindled C57BL/6J mice elicited Racine grade 4 seizures that were followed by PGES (Fig. 1A, B). The mean seizure duration was 36.64 ± 11.83 s. Seizures ranged from 13.60-75.00 s with most (71.58%) seizures lasting between 25-45 s (Fig. 1C). The mean PGES duration was 47.14 ± 36.26 s with a range between 8.75-233.10 s. Most (81.4%) mice exhibited PGES between 20-60 s (n = 55, 32 male, 23 female; Fig. 1D). There was no correlation between seizure duration and PGES duration, indicating that effects on PGES are not simply due to a decrease in seizure length (Fig. 1E).

**Figure 1.**
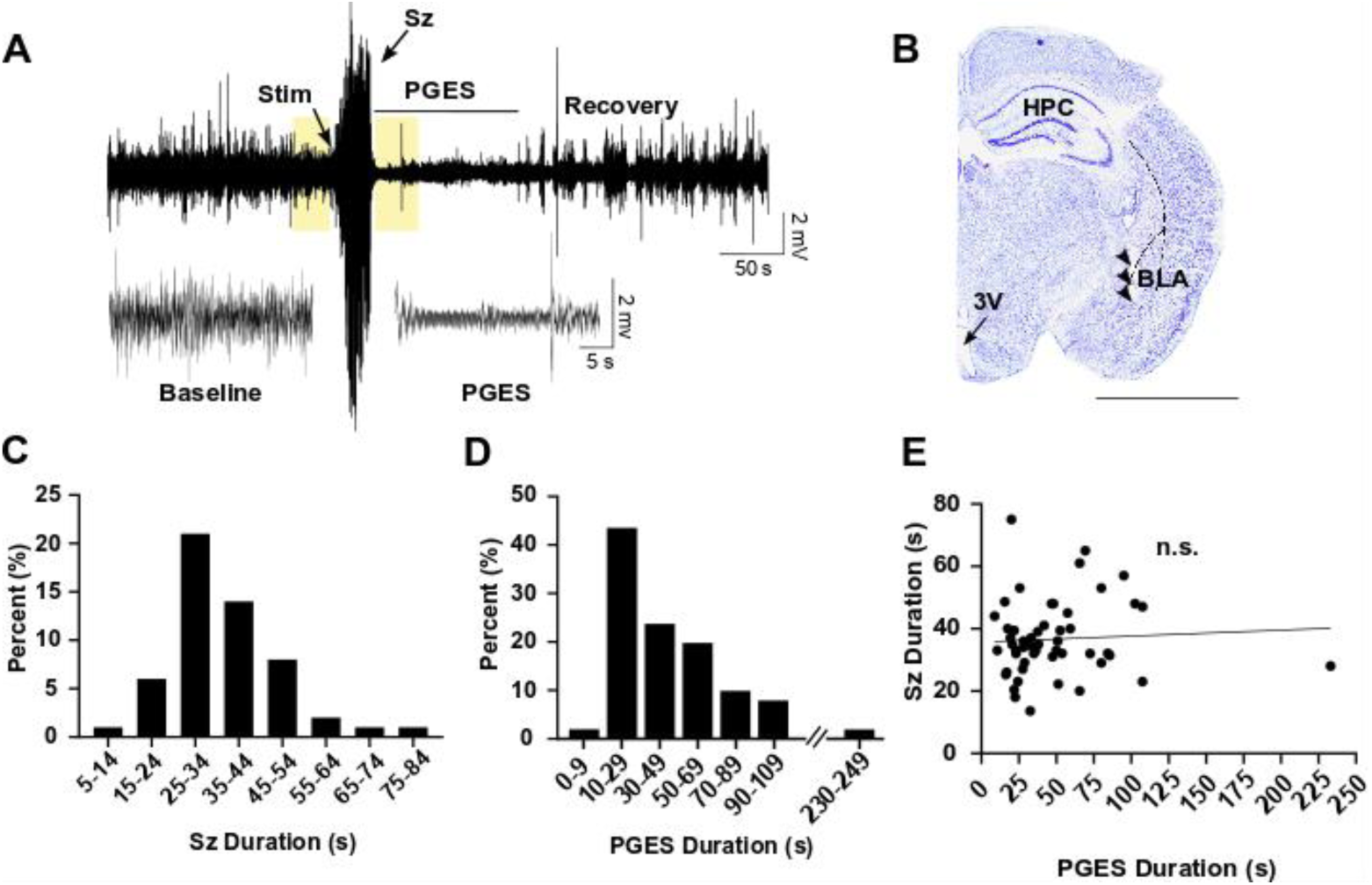
PGES was consistently observed following kindled seizures in amygdala kindled mice. (A) Top: 500 s raw EEG trace depicting seizure and PGES (duration indicated by horizontal line) in a C57BL/6 mouse. Scale bars: horizontal, 50 s; vertical, 2 μV. Bottom traces correspond to boxes on top trace and demonstrate EEG at baseline and during PGES. Scale bars: horizontal, 5 s; vertical 1 μV. (B) Nissl stained coronal hemi-section demonstrating electrode tract (arrowheads) in the basolateral amygdala. HPC, hippocampus; 3V, third ventricle; BLA, basolateral amygdala. Scale bar: 3 mm. (C,D) seizure durations (C) and PGES durations (D). (E) Linear regression analysis of seizure duration in amygdala kindled mice versus PGES duration.

### Systemic application of SSRIs decreased PGES duration following amygdala kindled seizures

To determine whether increasing 5-HT could reduce PGES duration, the SSRIs citalopram and fluoxetine, or vehicle, were systemically administered (*i*.*p*.) to mice prior to seizures induced during wake or NREM. Compared to saline treatment, citalopram (20 mg/kg) administration prior to seizure induction reduced PGES duration from seizures induced during wake (52.33 ± 8.336 s vs 18.54 ± 2.345 s; n = 30, 23; p < 0.05; Fig. 2A, E) and NREM sleep (37.16 ± 17.18 s vs 20.00 ± 7.84 s, n = 13; p < 0.05; Fig. 2A, F). Citalopram also reduced the duration of seizures induced during wake (38.43 ± 2.234 s vs 30.72 ± 2.704 s; n = 30, 23; p < 0.05; Fig. 2C), but not NREM sleep (34.59 ± 2.578 s vs 32.26 ± 3.274 s; n = 18, 14; Fig. 2C). Pretreatment with fluoxetine (10 mg/kg) significantly reduced PGES following seizures induced during wake (52.33 ± 45.66 s vs 28.25 ± 13.31 s, n = 11; p < 0.05; Fig. 2B, G), but not seizures induced during NREM (52.33 ± 8.336 s vs 27.97 ± 4.96 s; n = 30, 11; Fig. 2B, H). The duration of seizures induced during wake (38.43 ± 2.234 s vs 35.91 ± 2.04 s; n = 30, 11; Fig. 2D) and NREM (34.59 ± 2.57 s vs 36.22 ± 2.53 s; n = 18, 9; Fig. 2D) was unaffected by fluoxetine administration.

**Figure 2.**
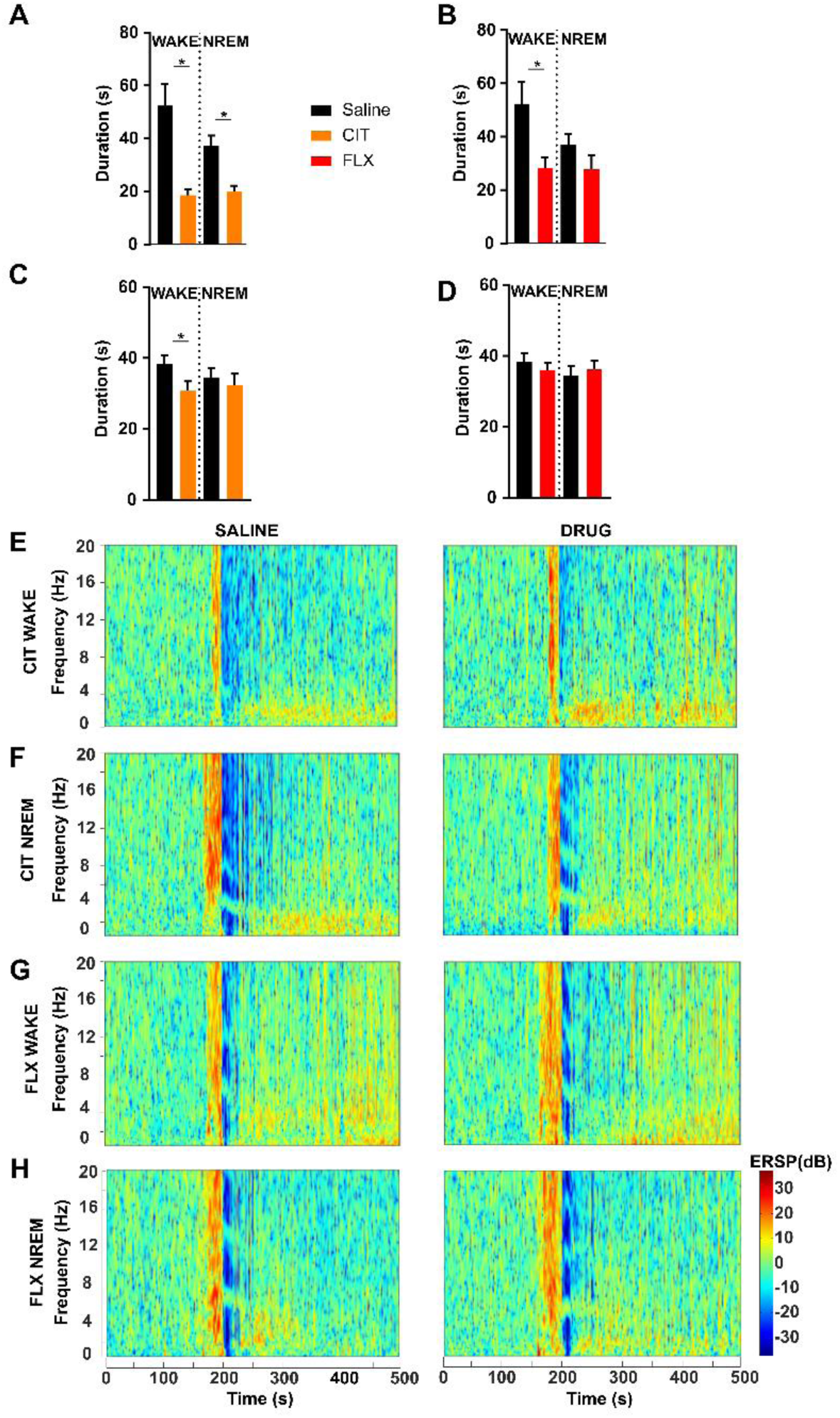
Systemic application of SSRIs reduced PGES following amygdala kindled seizures. (A,B) PGES duration and seizure duration (C,D) following seizures induced during wake and NREM in mice pre-treated with citalopram (A,C; CIT; 20 mg/kg, *i*.*p*.) or fluoxetine (B,D; FLX; 10 mg/kg, *i*.*p*.). *, p < 0.05. (E-H) Representative spectral plots demonstrating PGES duration following amygdala kindled seizures in mice that received pretreatment with (E,F) CIT or (G,H) FLX prior to seizures induced during wake (E,G) or NREM (F,H).

### Stimulation of DRN with acidosis reduced PGES following amygdala kindled seizures

To determine whether stimulation of DRN neurons prior to a seizure could reduce PGES duration, adult male and female C57BL/6J mice were instrumented with a guide cannula aimed at the DRN. 5-HT neurons within the DRN are chemosensitive and respond robustly to decreases in pH (Richerson, 2004), therefore, to stimulate DRN 5-HT neurons we focally administered normal or acidified aCSF through the guide cannula for 10 min prior to seizure induction. Pre-seizure perfusion of acidified aCSF to the DRN reduced PGES duration following seizures induced during wake (43.75 ± 6.73 s vs 25.79 ± 4.95 s; n = 6; p < 0.05; paired t-test; Fig. 3A, D). There was no effect on seizure duration (Fig. 3B). Application of acidified aCSF to the DRN in this manner during NREM sleep causes arousal (Smith, et al., 2018), so the effect of DRN stimulation on seizures induced during sleep could not be tested.

**Figure 3.**
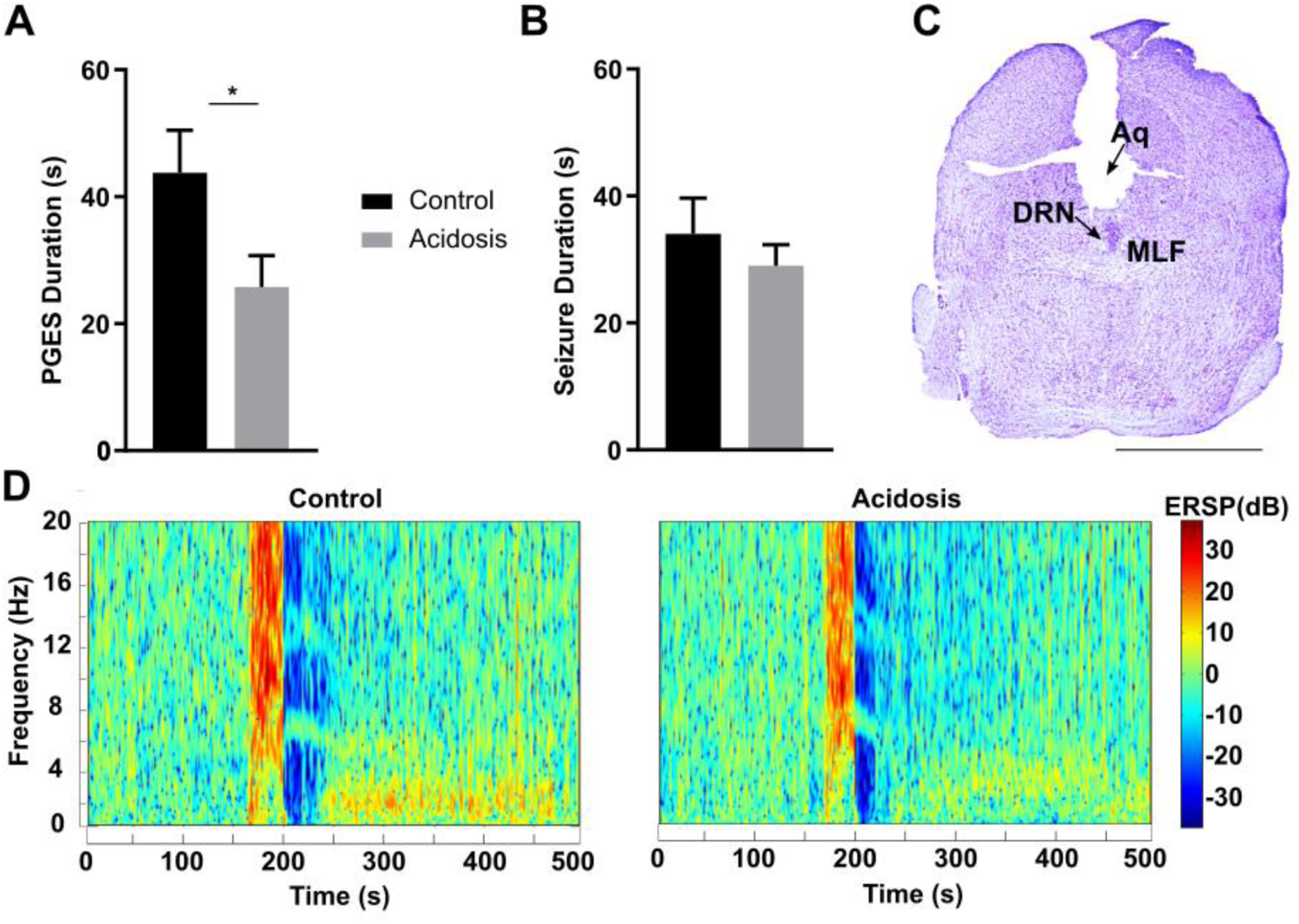
Focal chemical stimulation of the DRN reduced PGES following amygdala kindled seizures. (A,B) PGES duration (A) and seizure duration (B) following perfusion of normal (pH 7.4) or acidified (pH 6.8) aCSF prior to seizure induction during wake. *, p < 0.05. (C) Nissl stained coronal section at the level of the midbrain depicting position of the cannula targeted toward the DRN. Aq, aqueduct; DRN, dorsal raphe nucleus; MLF, medial longitudinal fasciculus; scale bar: 2.5 mm. (D) Representative spectral plots of PGES duration following perfusion of normal (left) or acidified (right) aCSF prior to seizure induction.

### Stimulation of DRN with acidosis reduced mortality following MES seizures

While stimulation of DRN with acidosis reduced PGES duration following seizures induced by amygdala stimulation in amygdala kindled mice, there is not mortality in the model. To determine whether stimulation of DRN neurons with acidosis could also reduce mortality, DRN neurons were stimulated with acidified aCSF for 10 min prior to seizure induction via MES, a model with known mortality (Buchanan, et al., 2014; Hajek and Buchanan, 2016; Kruse et al., 2019; Li and Buchanan, 2019; Purnell, et al., 2017). Perfusion of acidified aCSF into the DRN prior to MES induced seizures reduced mortality (4 survived vs 2 died; acidosis: n = 6; normal: n = 6; p < 0.05; Fig. 4A). Application of acidified aCSF to the DRN had no effect on seizure duration (Fig. 4B) and seizure duration did not differ between animals that survived versus died (independent t-test) (Fig. 4C). Seizure severity was similarly unaffected and did not differ between animals that survived versus died (Fig. 4D, E).

**Figure 4.**
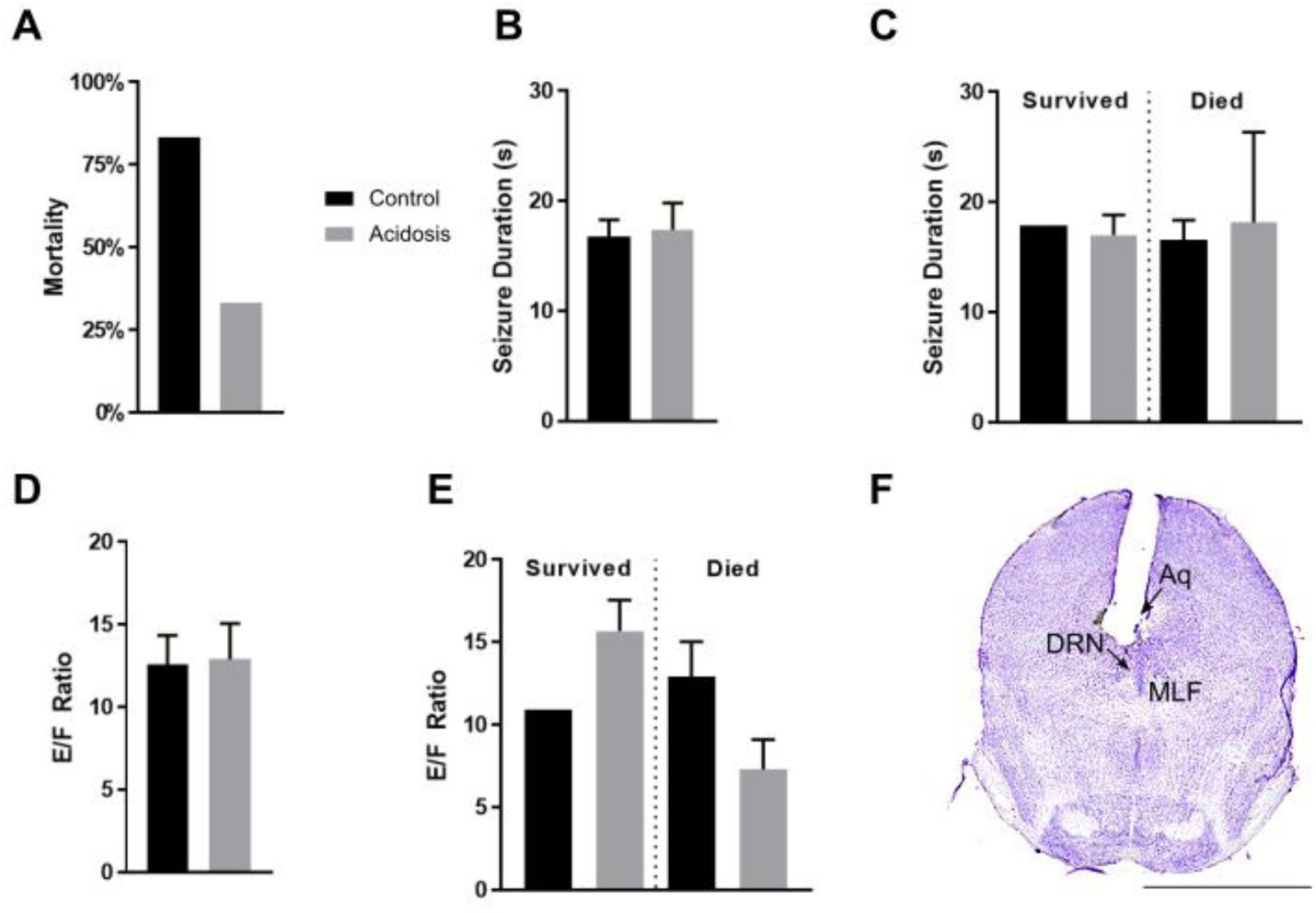
Focal chemical stimulation of the DRN reduced mortality following MES seizures without affecting seizure duration or severity. (A,B) Seizure-induced mortality (A) and seizure duration (B) from seizures induced following perfusion of normal (pH 7.4) or acidified aCSF (pH 6.8) into the DRN. (C) Seizure duration in mice that survived vs died from the MES seizure. (D) Seizure severity (E/F ratio) following pre-seizure perfusion of acidified aCSF. (E) Seizure severity in mice that survived vs died from the MES seizure. (F) Nissl stained coronal section at the level of the midbrain demonstrating cannula targeted toward the DRN. Aq, aqueduct; DRN, dorsal raphe; MLF, medial longitudinal fasciculus; scale bar: 2.5 mm.

### Optogenetic stimulation of DRN 5-HT neurons reduced PGES following amygdala kindled seizures

Kindled *Pet1-Cre* or *WT* mice that were injected with AAV-ChR2 into the DRN were subjected to laser light stimulation or no laser light stimulation for 10 min prior to induction of an amygdala kindled seizure during wake. Application of laser light to DRN 5-HT neurons prior to seizure induction during wake decreased PGES (100.4 ± 26.82 s vs 37.57 ± 13.22 s; n = 8; p < 0.05; Wilcoxon matched pairs signed rank test; Fig. 5A, F) without affecting seizure duration (*Pet1-Cre*: 29.57 ± 5.54 s vs 29.61 ± 2.83 s; n = 8; *WT*: 22.14 ± 1.895 s vs 23.14 ± 1.62 s; n = 7; Fig. 5B). Viral transfection in *Pet1-Cre* mice was successful and limited to our 5-HT cell populations of interest (Fig. 5C-E). PGES duration and seizure duration did not differ between *Pet1-Cre* mice and their *WT* littermates (Fig. 5A, B, F). There was no ChR2 expression in *WT* mice (Fig. 5E).

**Figure 5.**
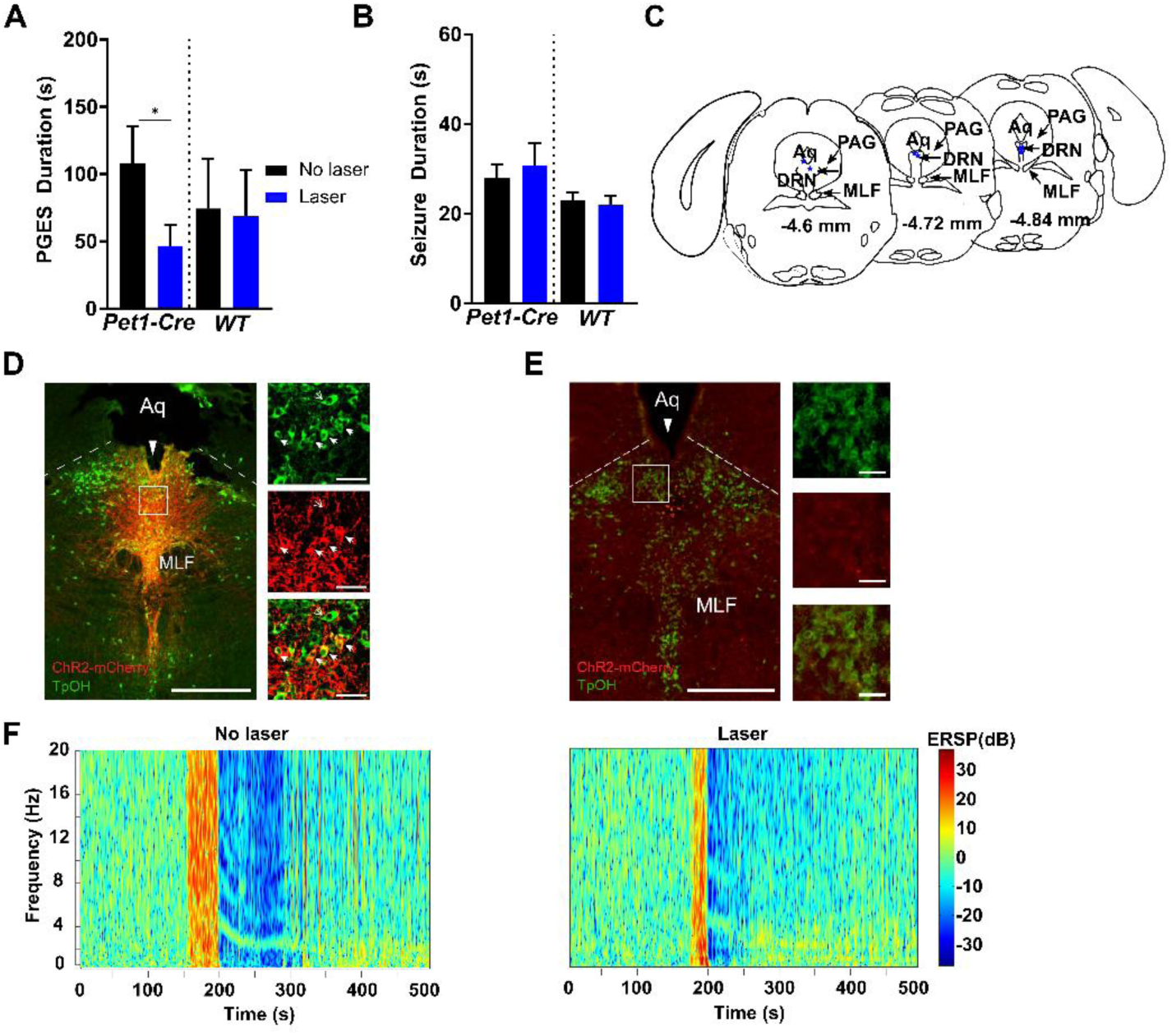
Optogenetic stimulation of the DRN reduced PGES following amygdala kindled seizures. (A, B) PGES duration (A) and seizure duration (B) following laser stimulation (473 nm, blue; 4 Hz, 10 mW, 2% duty; 10 s ON / 10 s OFF; ChR2: n = 7; *WT*: n = 8) or no stimulation (black) of DRN neurons prior to seizure induction during wake. *, p < 0.05. (C) Depiction of optic fiber positioning within the DRN in *Pet1-Cre* mice injected with AAV-ChR2. Aq, aqueduct; DRN, dorsal raphe nucleus; PAG, periaqueductal gray; MLF, medial longitudinal fasciculus. (D,E) Immunostained coronal section depicting optical fiber tracts (long solid arrow), ChR2 expression (red), 5-HT cell bodies (green/non-solid small arrow), and co-expression (yellow/solid small arrows) within the DRN of a *Pet1-Cre* (D) and *WT* (E) mouse that received an injection of AAV-ChR2 into DRN. Box indicates location of magnified cells on right (top, 5-HT; middle, ChR2; bottom, merge). Aq, aqueduct; MLF, medial longitudinal fasciculus; scale bar: 500 µm, 50 µm. (F) Representative spectral plots of PGES duration following either no laser (left) or laser stimulation (right) of DRN 5-HT neurons prior to seizure induction in a *Pet1-Cre* mouse injected with AAV-ChR2.

### PGES in amygdala kindled mice was associated with immobility that could be prevented by pretreatment with citalopram

During PGES in amygdala kindled mice, there was immobility that dissipated in a stepwise manner. Mice were immobile at the onset of PGES. This gradually progressed sequentially to a body twitch, then unilateral paw movements, and then finally bilateral effort to reposition the body or resumption of preening behavior (n = 55; Fig. 6A). The stepwise return of mobility was observed during PGES following seizures induced during wake (twitch: 15.08 ± 1.81 s vs paw movement: 28.40 ± 2.89 s vs body movement: 51.27 ± 5.95 s; n = 31; p < 0.05) and NREM sleep (twitch: 16.78 ± 2.52 s vs paw movement: 29.69 ± 4.74 s vs body movement: 60.17 ± 9.60 s; n = 18; p < 0.05) with no difference in the latency to each phase (Fig. 6B).

**Figure 6.**
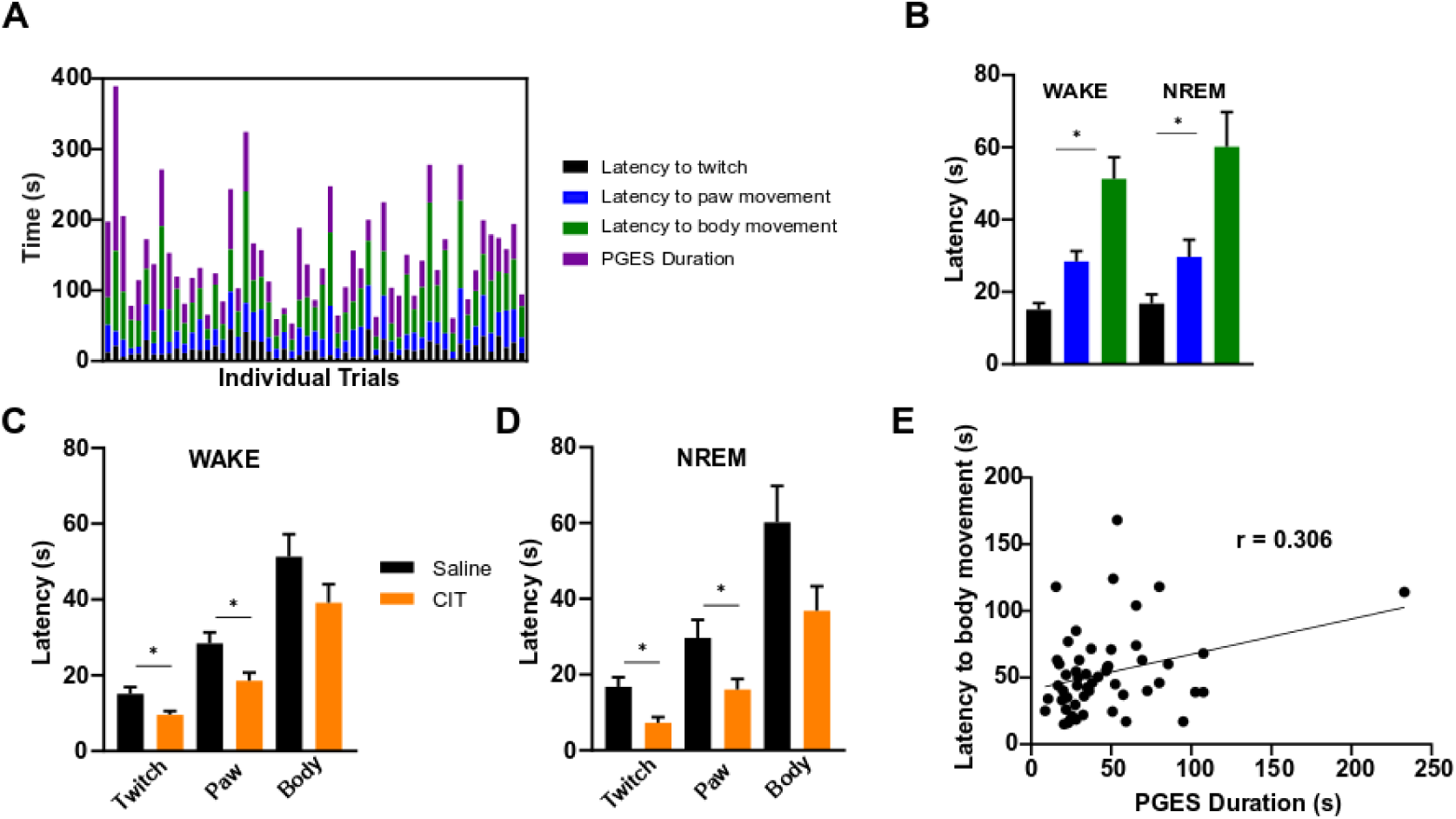
PGES was associated with immobility that could be reversed by administration of citalopram. (A) Latency to observation of first twitch (black), paw movement (blue), and body movement (green) after end of seizure. Duration of PGES (purple) is plotted for each trial. Columns depict a different animal/trial (n = 55). (B) Mean latencies to first observed twitch (black), paw movement (blue), and body movement (green) after end of seizure induced during wake (n =31) and NREM (n = 19). *, p < 0.05. (C,D) Latency to observation of first twitch, paw movement, and body movement after end of seizure induced during wake (C; n = 23) and NREM (D; n = 13) in mice pretreated with saline (black) or CIT (orange; 20 mg/kg, *i*.*p*.). *, p < 0.05. (E) Linear regression between PGES duration and latency to body movement following seizures induced during wake (n = 31). *, p < 0.05.

**Figure 7.**
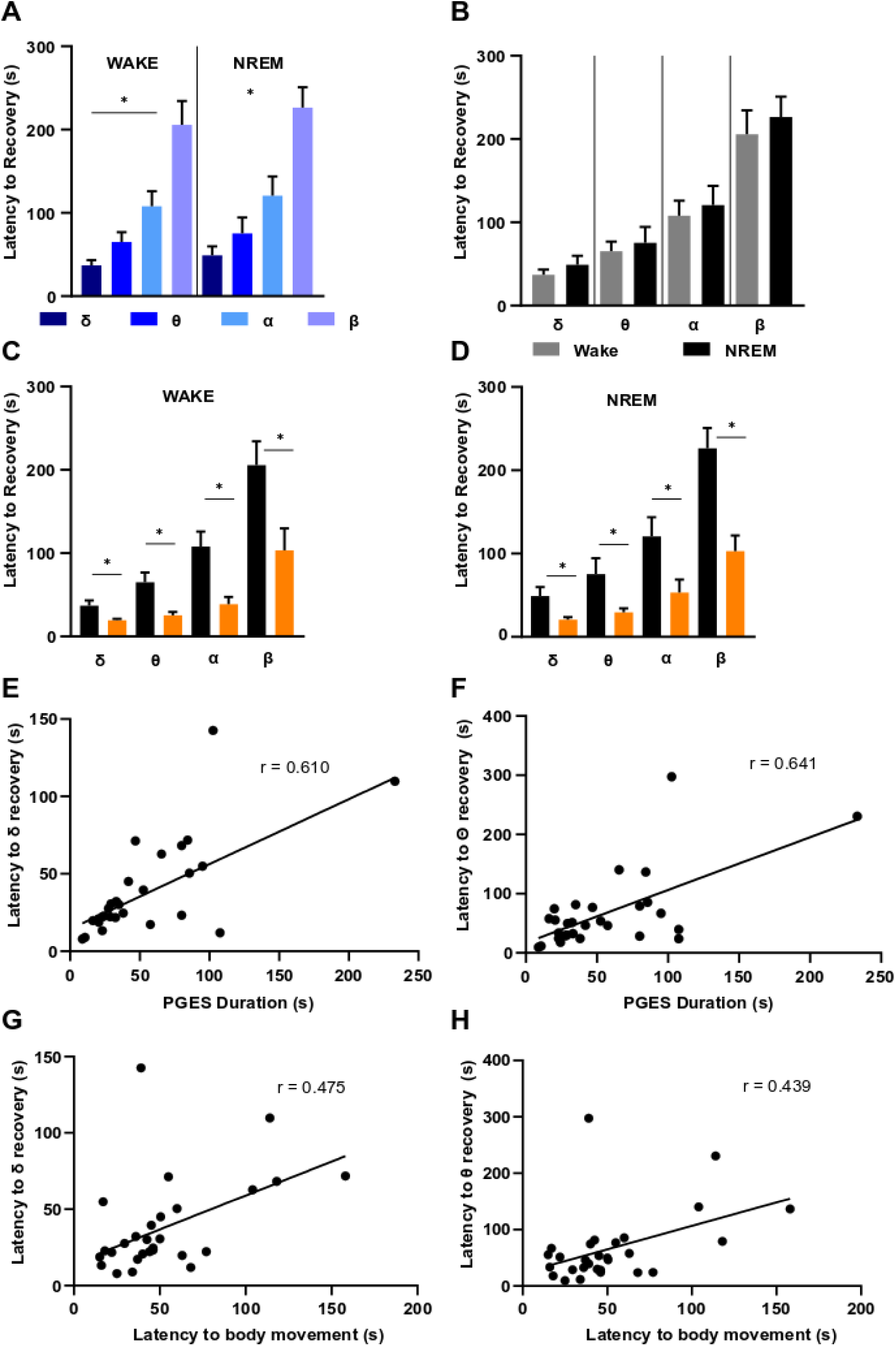
Resumption of baseline EEG activity occurred in a stepwise manner during PGES and could be accelerated with citalopram pretreatment. (A,B) Recovery of EEG rhythms to baseline during PGES following seizures induced during (A) wake (δ, θ, α: n = 31; β: wake: n = 26) and (B) NREM (δ, θ, α: n = 19; β: n = 13). *, p < 0.05. (C,D) Effect of citalopram pretreatment (CIT; *i*.*p*., 20 mg/kg) 30-60 min before seizure induction during (C) wake (δ, θ, α: n = 23; β: n = 12) and (D) NREM (δ, θ, α: n = 13, 8 male, 5 female; β: n = 7, 4 male, 3 female; Mann-Whitney U test) on recovery of EEG rhythms to baseline during PGES. *, p < 0.05. (E,F) Linear regression analysis of the relationship between recovery of δ (E, r = 0.610) and θ (F, r = 0.641) with PGES duration following seizures induced during wake. *, p < 0.05. (G,H) Linear regression of the relationship between recovery of δ (G, r = 0.475) and θ (H, r = 0.439) with latency to body movement during PGES. *, p < 0.05. delta (δ), 1 – 4 Hz; theta (θ), 4 – 8 Hz; alpha (α), 8 – 12 Hz.

Pretreatment with citalopram (*i*.*p*., 20 mg/kg) 30-60 min before seizure induction during wake (twitch: 15.08 ± 1.81 s vs 9.591 ± 4.56 s; paw movement: 28.40 ± 2.89 s vs 18.55 ± 10.11 s; body movement: 51.27 ± 5.94 s vs 39.19 ± 2.73 s; n = 31, 22; p < 0.05) and NREM (twitch: 16.78 ± 2.52 s vs 7.27 ± 1.48 s; paw movement: 29.69 ± 4.74 s vs 16.00 ± 2.85 s; body movement: 60.17 ± 9.60 s vs 36.82 ± 6.49 s; n = 18, 11; p < 0.05) sleep reduced the latency to first twitch and paw movement (Fig. 6C, D). Fluoxetine administration prior to seizures induced during wake (twitch: 15.08 ± 1.81 s vs 11.91 ± 1.80 s; paw movement: 28.40 ± 2.89 s vs 25.50 ± 3.43 s; body movement: 51.27 ± 5.94 s vs 48.48 ± 6.31 s; n = 31, 11) and NREM (twitch: 16.78 ± 2.52 s vs 15.28 ± 2.96 s; paw movement: 29.69 ± 4.74 s vs 35.61 ± 6.80 s; body movement: 60.17 ± 9.60 s vs 52.00 ± 7.23 s; n = 18, 11) did not reduce latency to twitch, paw movement, or body movement during PGES. There was a positive correlation observed between latency to body movement and PGES duration in seizures induced during wake (n = 31; p < 0.05; r = 0.306; Fig. 6E), but not NREM (n = 18; r = 0.09).

Pretreatment with citalopram (*i*.*p*., 20 mg/kg) 30-60 min before seizure induction during wake (twitch: 15.08 ± 1.81 s vs 9.591 ± 4.56 s; paw movement: 28.40 ± 2.89 s vs 18.55 ± 10.11 s; body movement: 51.27 ± 5.94 s vs 39.19 ± 2.73 s; n = 31, 22; p < 0.05) and NREM (twitch: 16.78 ± 2.52 s vs 7.27 ± 1.48 s; paw movement: 29.69 ± 4.74 s vs 16.00 ± 2.85 s; body movement: 60.17 ± 9.60 s vs 36.82 ± 6.49 s; n = 18, 11; p < 0.05) sleep reduced the latency to first twitch and paw movement (Fig. 6C, D). Fluoxetine administration prior to seizures induced during wake (twitch: 15.08 ± 1.81 s vs 11.91 ± 1.80 s; paw movement: 28.40 ± 2.89 s vs 25.50 ± 3.43 s; body movement: 51.27 ± 5.94 s vs 48.48 ± 6.31 s; n = 31, 11) and NREM (twitch: 16.78 ± 2.52 s vs 15.28 ± 2.96 s; paw movement: 29.69 ± 4.74 s vs 35.61 ± 6.80 s; body movement: 60.17 ± 9.60 s vs 52.00 ± 7.23 s; n = 18, 11) did not reduce latency to twitch, paw movement, or body movement during PGES. There was a positive correlation observed between latency to body movement and PGES duration in seizures induced during wake (n = 31; p < 0.05; r = 0.306; Fig. 6E), but not NREM (n = 18; r = 0.09).

### Citalopram reduced latency to return of EEG activity to baseline during PGES

EEG bands re-emerged sequentially during PGES with the slower, low frequency bands recovering before the high frequency bands (Fig. 7A). This pattern was observed during PGES following seizures induced during wake (δ: 38.24 ± 31.20 s; θ: 67.60 ± 63.02 s; α: 102.70 ± 82.43 s; β: 183.30 ± 104.50 s; p < 0.05) or NREM (δ: 46.37 ± 42.75 s; θ: 73.41 ± 74.76 s; α: 115.1 ± 91.53 s; β: 223.60 ± 78.13 s; p < 0.05) sleep (Fig. 7B). Pretreatment with citalopram (*i*.*p*., 20 mg/kg) 30-60 min before seizure induction during wake (δ: 18.27 ± 7.60 s; θ: 24.25 ± 14.15 s; α: 37.50 ± 36.63 s; β: 87.80 ± 81.49 s; δ, θ, α, wake: n = 23; β, wake: n = 12; p < 0.05) and NREM (δ: 20.25 ± 9.613 s; θ: 28.09 ± 15.50 s; α: 50.03 ± 53.68 s; β: 102.9 ± 49.37 s; δ, θ, α, NREM: n = 13; β, NREM: n = 7; p < 0.05) sleep reduced the latency of all frequencies to return to baseline activity (Fig. 7C, D). Pretreatment with fluoxetine did not affect the latency of EEG rhythms returning to baseline activity following seizures induced during wake, but did following NREM (δ: 31.63 ± 3.73 s; θ: 39.28 ± 5.92 s; α: 73.34 ± 15.58 s; β: 148.3 ± 23.78 s; δ, θ, α, β wake: n = 11; δ: 24.47 ± 5.33 s; θ: 33.05 ± 4.49 s; α: 59.72 ± 20.73 s; β: 106.2 ± 32.57 s; NREM: n = 9; p < 0.05). Latency to baseline activity of the δ and θ frequency bands correlated most strongly with PGES duration in seizures induced during wake (δ: r = 0.610; θ: r = 0.641; α: r = 0.442; β: r = 0.478; p < 0.05). Similarly, latency to full body movement also correlated with return to baseline activity of the δ and θ frequency bands in seizures induced from wake (δ: r = 0.47; θ: r = 0.44; p < 0.05). However, these correlations were not observed for seizures induced from NREM (δ: r = 0.044; θ: r = 0.094).

## Discussion

Prolonged PGES duration may portend increased risk of SUDEP. Understanding methods of shortening PGES duration may translate to methods to reduce the likelihood of SUDEP. However, mechanisms underlying PGES and thus ways to shorten it, are unknown. Here we found that stimulating serotonergic neurotransmission at the DRN reduced PGES duration and could also reduce seizure-related mortality.

PGES is common following seizures in humans. It is most common with generalized seizures (Surges et al., 2011), whether they are primary generalized (Poh et al., 2012) or focal to bilateral tonic clonic seizures (Marchi et al., 2019), but can also be seen with focal seizures (Lhatoo, et al., 2010). Similarly, PGES has been observed in animal models following kindled seizures (Buterbaugh, 1987), MES seizures (Hajek and Buchanan, 2016), and pentylenetetrazol induced seizures (Lüttjohann et al., 2009; Mirski and Fisher, 1994). PGES preceded all cases with adequate data in the multi-center MORTEMUS study. Furthermore, early research in humans suggests that PGES duration exceeding 50 s increases SUDEP risk (Lhatoo, et al., 2010).

PGES often follows nocturnal (Alexandre et al., 2015; Latreille et al., 2017; Okanari, et al., 2017) and generalized tonic clonic seizures (Lhatoo, et al., 2010; Surges, et al., 2011), which are independent risk factors for SUDEP (Lamberts, et al., 2012). PGES has also been associated with peri-ictal tachycardia and hypoxemia in pediatric patients (Moseley et al., 2013). Increases in sympathetic tone and decreases in parasympathetic tone observed during PGES may also precipitate fatal cardiac arrhythmias (Bozorgi et al., 2013; Moseley et al., 2013; Poh, et al., 2012). In turn, cardiac dysfunction may further precipitate PGES by reducing blood flow and oxygenation to the brain (Bozorgi and Lhatoo, 2013).

It should be noted that while some studies observed a positive correlation between PGES and SUDEP (Lhatoo, et al., 2010), other studies have not recapitulated this finding (Lamberts et al., 2013; Surges, et al., 2011). Surges et al. observed no difference in the presence or duration of PGES between SUDEP and control cases (Surges, et al., 2011). Other studies have not supported the observation that PGES is more likely to follow convulsive seizures (Lamberts, et al., 2013; Xu et al., 2016). Moreover, despite suppression of scalp EEG activity, subcortical activity may persist during PGES (Bateman et al., 2019). However, another study indicated that motor artifact (oral tonicity, facial automatisms) and proximity of electrodes to the skull can cause detection of gamma activity during PGES (Marchi, et al., 2019; McGonigal et al., 2019; Okanari, et al., 2017). It is possible that small movements may have contributed to their findings, but it is also likely that not all brain regions are equally suppressed during PGES. Faster recovery may occur in subcortical structures that are beyond detection on scalp EEG. Continued PGES research in animal models may provide insight into such questions that highly variable human studies cannot.

Additionally, impairment of arousal is relevant to PGES. SUDEP victims are often found deceased in a prone position with an obstructed airway (Kloster and Engelskjon, 1999; Liebenthal et al., 2015; Nashef et al., 2007; Tao et al., 2010; Zhuo et al., 2012). Seizures with PGES exhibit longer post-ictal immobility (Asadollahi et al., 2018; Seyal et al., 2013) which may increase risk of rebreathing CO_2_ and exacerbate unconsciousness. This immobility was recapitulated in our mice. Mice progressively recovered mobility (twitch → paw movement → body movement) following seizures. Pretreatment with citalopram not only shortened PGES duration, but shortened latency to first twitch and paw movement in our mice. It is difficult to correlate post-ictal movement in a mouse to an epilepsy patient; however, it may provide an adequate approximation for when life-saving responses to stimuli may reoccur.

Reduced arousal, stupor, and unconsciousness are frequently associated with generalized seizures (Blumenfeld, 2012; Englot et al., 2010) and are associated with PGES (Kuo, et al., 2016). An inability to arouse to CO_2_ or other stimuli during the postictal period may be one mechanism by which PGES contributes to seizure mortality. We hypothesize that PGES may be an electrographic marker of impaired arousal consequent to ictal 5-HT neurotransmission dysfunction. Here administration of SSRIs prior to a seizure decreased PGES duration in kindled mice. Similarly, pre-seizure application of acidified aCSF to the DRN or optogenetic stimulation of DRN 5-HT neurons decreased PGES duration in kindled mice. In the MES model, PGES duration and mortality is greater following seizures induced during NREM sleep (Hajek and Buchanan, 2016; Purnell, et al., 2017). Our results indicate that acidosis of the DRN reduced mortality following MES seizures induced from wake. Application of acidified aCSF to the DRN precluded induction of seizures during sleep. However, it is likely that seizure induction during NREM/REM may have increased mortality despite DRN acidosis due to decreases in 5-HT activity during sleep (McGinty and Harper, 1976). It is possible that DRN acidosis reduced mortality by preventing propagation of the seizure to the DRN or reducing the effect of seizures on DRN activity. We did not observe an increase in PGES following kindled seizures induced in NREM; however, this may be due to differing seizure propagation and affected regions between the MES and kindling model.

In humans, PGES is often followed by δ slowing (Fisher and Engel, 2010; Kaibara and Blume, 1988) that eventually transitions into θ as the EEG begins to return to normal (Fisher and Engel, 2010; So and Blume, 2010). PGES duration correlated most strongly with the recovery latency of the δ and θ bands. Cortical δ slow waves are often associated with reduced consciousness and responsiveness to stimuli (Blumenfeld, 2012). Disruption of brainstem ascending arousal system structures by seizures may lead to depressed cortical activity and detection of cortical slow waves (Blumenfeld, 2012). It is possible that the emergence of slow waves following PGES indicates resumption of some subcortical activity, but a continued suppression of cortical activity. The DRN is a major component of the ascending arousal system that sends 5-HT projections to the cortex (Vertes, 1991). Stimulation of the DRN with acidosis may have reduced PGES by increasing activity of its projections to the cortex, thus restoring reciprocal interactions between cortical-subcortical networks and terminating PGES.

The θ rhythm is primarily generated in the hippocampus (Adamantidis et al., 2019; Lubenov and Siapas, 2009) and is driven by GABAergic neurons in the medial septum firing at 4-8 Hz (Colgin, 2016). It is plausible that as EEG recovery continues following PGES, the emergence of θ waves may be an EEG indicator of continued post-ictal stupor and confusion. Persistence of the θ rhythm following waking is associated with disorientation, reduced vigilance, and reduced arousal in humans (Ferrara et al., 2006). We also observed a correlation between the recovery of the δ and θ frequency bands and latency to full body movement in our mice. During wakefulness, hippocampal θ activity is associated with locomotion, spatial navigation, and locomotion speed (Bender et al., 2015; Slawinska and Kasicki, 1998). Therefore, increases in θ activity may have preceded or followed movement initiated by the mice as they recovered from PGES.

To conclude, we do not understand the origin, significance, or mechanisms producing PGES. Very few studies have addressed PGES in animal models. Here we demonstrate that PGES occurs consistently following seizures in kindled animals and that increasing 5-HT activity can reduce PGES duration, mortality, and immobility following PGES. PGES may result from seizure-induced depression of subcortical arousal network activity to the cortex. The DRN may be integral because seizures reduce firing of DRN 5-HT neurons (Zhan, et al., 2016) and, under normal conditions, the DRN sends 5-HT projections to the hippocampus, the pedunculopontine tegmental nucleus, the basal forebrain, the cortex, and other wake promoting structures (Monti, 2010; Vertes, 1991). While the DRN is likely not the only structure involved in the generation of PGES, it is evident that increasing 5-HT activity prior to a seizure reduces PGES and post-ictal immobility. More work will be needed to understand the precise location and mechanism of action of 5-HT in modulating PGES. Ultimately, it is our hope that enhancing knowledge of PGES will aid in identifying high-risk epilepsy patients and reduce seizure-induced mortality.

## Acknowledgements

This work was supported by a Post-Comprehensive Exam Fellowship from the Graduate College at the University of Iowa and a Pre-Doctoral Fellowship from the American Epilepsy Society (with funding from LivaNova, PLC) to A.N.P., a Summer Fellowship from the Iowa Center for Research by Undergraduates to K.M.V., and NIH/NINDS R01 NS095842 and the Beth L. Tross Epilepsy Professorship from the Carver College of Medicine at the University of Iowa to G.F.B. Funding sources did not influence study design, data collection, analysis, interpretation, preparation of the manuscript, or the decision to publish.

## Conflict of interest statement

The authors declare no competing financial interests.

## Abbreviations

3V: third ventricle
5-HT: 5-hydroxytryptamine, serotonin
AAV: adeno-associated virus
aCSF: artificial cerebrospinal fluid
AP: anterior-posterior
Aq: aqueduct
BLA: basolateral amygdala
ChR2: channelrhodopsin2
CIT: citalopram
Cy: cyanine
DRN: dorsal raphe nucleus
Ds: discosoma
DV: dorsal-ventral
EEG: electroencephalogram
E/F ratio: extension-to-flexion ratio
EMG: electromyogram
ERSP: event-related spectral perturbation
FLX: fluoxetine
GEPR: genetic epilepsy prone rats
GFP: green fluorescent protein
HPC: hippocampus
MES: maximal electroshock
ML: medial-lateral
MLF: medial longitudinal fasciculus
NREM: non-rapid eye movement
PAG: periaqueductal gray
PBS: phosphate buffered saline
PGES: post-ictal generalized EEG suppression
SSRI: selective serotonin reuptake inhibitor
SUDEP: sudden unexpected death in epilepsy
Sz: seizure
TpOH: tryptophan hydroxylase
WT: wild type

## Contributions

A.N.P. and G.F.B. conceived and designed the research; A.N.P., K.G.J., J.W.C, and K.M.V. performed experiments; A.N.P., K.G.J., J.W.C., R.L., and G.F.B. analyzed data and interpreted results of experiments; A.N.P. prepared figures and drafted manuscript; A.N.P., K.G.J., J.W.C., R.L, K.M.V, and G.F.B. revised and approved final version of the manuscript.

## Notes

### Competing Interest Statement

The authors have declared no competing interest.

